# Precision motor timing via scalar input fluctuations

**DOI:** 10.1101/2022.05.18.492498

**Authors:** Rich Pang, Alison Duffy, David Bell, Zsofia Torok, Adrienne Fairhall

## Abstract

Complex motor skills like playing piano require precise timing over long periods, without errors accumulating between subprocesses like the left and right hand movements. While biological models can produce motor-like sequences, how the brain quenches timing errors is not well understood. Motivated by songbirds, where the left and right brain nuclei governing song sequences do not connect but may receive low-dimensional thalamic input, we present a model where timing errors in an autonomous sequence generator are continually corrected by one-dimensional input fluctuations. We show in a spiking neural network model how such input can rapidly correct temporal offsets in a propagating spike pulse, recapitulating the precise timing seen in songbird brains. In a reduced, more general model, we show that such timing correction emerges when the spatial profile of the input over the sequence sufficiently reflects its temporal fluctuations, yielding *time-locking attractors* that slow advanced sequences and hasten lagging ones, up to the input timescale. Unlike models without fluctuating input, our model predicts anti-correlated durations of adjacent segments of the output sequence, which we verify in recorded zebra finch songs. This work provides a bioplausible picture of how temporal precision could arise in extended motor sequences and generally how low-dimensional input could continuously coordinate time-varying output signals.

**Significance:** Complex motor skills like playing piano require precision timing over long periods, often among multiple components like left and right muscle groups. Although brain-like network models can produce motor-like outputs, timing regulation is not well understood. We introduce a model, inspired by songbird brains, where imprecise timing in a cortical-like system is corrected by a single thalamic input regulating the sequential propagation, or tempo, of cortical activity. This model illuminates a relation between the input’s spatial structure and temporal variation that lets lagging activity hasten and advanced activity slow, which makes a prediction about output timing that we verify in real birdsong. This work reveals a simple, neuroplausible mechanism that may play a role in precision cortical or motor timing.

---

Complex motor sequences like playing the piano often require precision timing over extended periods. If small timing errors accumulated over long sequences, this could lead to substantial variability in sequence duration or desynchronize different subprocesses like a sequence’s left and right motor components, if their errors accumulated independently. While robust sequence generation by biological neural network models has been studied extensively (1–9), how the nervous system could prevent the accumulation of timing errors remains largely unexplored.

Brain regions associated with sequence generation do not operate in isolation but receive input from other areas. Mammalian motor cortex receives ongoing input from thalamus during movement, which if inhibited disrupts cortical patterning and limb kinematics (10). Feed-forward thalamic inputs are also implicated in preserving temporal information in sensory pathways (11). Although external inputs to a sequence-generating network may play multiple roles, such as initiating cortical state or movement (12–15), or gating signal transmission (16), one role relevant to timing control is modulating the speed at which the sequence-generating network’s dynamics unfold (17). It was shown that the level of external input to a recurrent neural network could adjust the interval from task start to output by smoothly scaling the individual response time-courses of its internal units (17). If fluctuating external input slowed or accelerated a network’s dynamics opposite their internal noisy temporal variations this might compensate for ongoing timing errors.

One of the most well-studied precision motor sequences is birdsong. Adult zebra finch song, a highly stereotyped vocalization sequence, lasts up to 2 seconds but varies only about 1.5% in total duration across renditions (18); other birds like canaries can sing for tens of seconds (19). Accompanying zebra finch song is a sparse, extremely precise spike sequence in premotor area HVC (proper name); many neurons burst during every song within a few-millisecond window surrounding a single song timepoint (20). While HVC is thought to be the central sequence generator underlying song timing (6), it requires input from the thalamic Uvaeform nucleus (Uva), which if lesioned abolishes or substantially distorts song (21). Uva activity is dynamic during song but highly correlated across the nucleus (21), suggesting it does not bequeath to HVC the complete sequential information guiding song but acts instead a global modulation signal, although its precise role is unknown. As songbirds have no corpus callosum, left and right HVC do not directly communicate (22); Uva inputs could in principle help them remain coordinated throughout song by dynamically controlling how fast each HVC’s spike sequence unfolds.

Here we introduce a model in which timing in an autonomous sequence generator is corrected purely via a onedimensional external input, inspired by Uva, that dynamically modulates the sequence’s propagation speed. In an HVC-like model network, we show how this input corrects timing errors in a propagating spike pulse even without access to the errors. Formulating the problem more generally we show this error correction occurs when the 1-D input is spread nonuniformly over the sequence so as to reflect the input’s time-derivative spatially, which yields “time-locking attractors” (fixed-point attractors (23) in a constant-velocity moving reference frame) that slow advanced sequences and hasten lagging ones. A key prediction of this model is an anti-correlation between durations of adjacent output segments, which we confirm in recorded zebra finch songs.

## Results

### Timing correction in a biological network model

To illustrate our proposed timing control mechanism we modeled a chain-organized neural network supporting the stable propagation of a spike pulse, in line with previous HVC models (5, 6, 24), but additionally modulated by an external 1-D input (Fig 1A). The chain comprised small recurrent clusters of adaptive spiking neurons connected via feed-forward excitatory synapses, with additional connections from the external input covering the whole chain, although nonuniformly. When the first cluster, or “link”, is stimulated, a brief spike pulse emerges that travels down the chain, which we take as our output sequence. Given that we will only consider sequences of a fixed order, the network state can be characterized during pulse propagation by *x*(*t*), the index (or “position”) of the sequence element active at time *t* (i.e. the chain link whose neurons are spiking at *t*) (Fig 1A-C). A spike pulse propagating at a constant speed, for example, would be described by *x*(*t*) = *vt*.

**Fig. 1.**
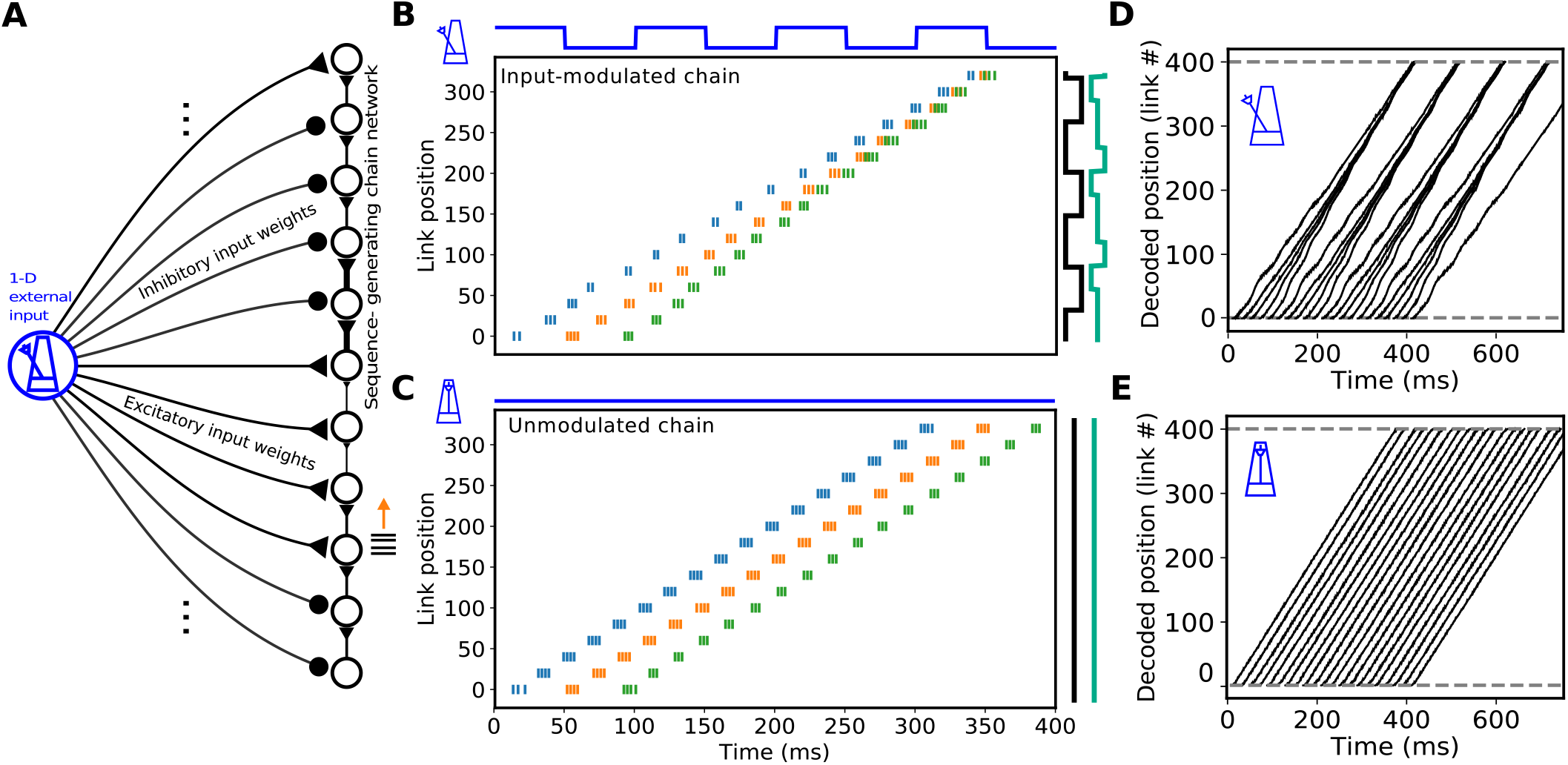
Timing correction via 1-D input in a simple spiking network model. A. Network architecture: A chain-organized network supporting the propagation of a spike pulse (right) is modulated by a 1-D input signal (blue) via either excitatory (triangles) or inhibitory (circles) weights that fall on distinct zones of the chain. When the input is on spike pulses propagate more quickly through excited zones and more slowly through inhibited zones. Momentary propagation speed is determined by the absence/presence and sign of the input and by the intrinsic chain speed governed by the feed-forward weights. B. Evolution of spike pulses in three separate trials, initiated at different start times in the input-modulated network. Only spikes from an evenly spaced subset of neurons are shown, labeled by the neurons’ chain link. Blue line shows input timecourse, black contour shows spatial modulation profile (left-inhibition, right-excitation), cyan contour shows three levels of intrinsic speed variation as a function of link position (slow, medium, fast from left to right). C. Propagation of three spike pulses in an unmodulated chain network with constant intrinsic speed. D. Evolution of spike pulse position (decoded from link positions of neurons spiking in a 2 ms window surrounding time point *t*) for a range of start times in the modulated network. E. As in D, but for the unmodulated network.

The propagation speed of the spike pulse when passing over a given link in the chain is determined by (1) the “intrinsic” chain speed (governed by the feed-forward connection strengths to and from that link), (2) the input weight at that link, and (3) the (scalar) activation level of the input when the pulse is passing over that link. Strong feedforward weights and active excitatory input hasten propagation; weak feed-forward weights and active inhibitory input slow propagation. Active excitatory input can also cancel the speed decrease from weak feed-forward weights, and active inhibitory input can cancel the speed increase from strong feed-forward weights. Here we assume a binary external input level (on or off) oscillating at 10 Hz, in line with observed Uva rhythms (21), although neither the specific frequency nor periodicity are essential to our mechanism. Because the input is 1-D, at any given time external inputs to the chain are all active or all silent, reflecting the putatitve low-dimensional nature of Uva (21). We let the spatial input profile alternate between excitation and inhibition and the intrinsic speed profile between weak, medium, and strong feed-forward weights (Fig 1A). For simplicity we modeled our Uva-inspired input as directly exciting or inhibiting different parts of the chain, but real Uva-HVC projections could be purely excitatory, although this is not yet known. It may thus be more realistic that any effective Uva-HVC inhibitory modulation is mediated through Uva projections to inhibitory interneurons in HVC, which in turn project to HVC excitatory neurons (25).

Given this network configuration we sought conditions under which large offsets in spike pulse initiation times were corrected by fluctuating input. An example solution is shown in Fig 1B. Given a 100 ms temporal modulation period and appropriate spatial variation (discussed shortly) in both the feed-forward weights in the chain and the input weight profile, spike pulses initialized either 40 ms before or after a “correct” start time cleanly hastened or slowed, respectively, after just a few hundred ms to approximately match the timing of the correct pulse. In an unmodulated chain network with spatially constant feed-forward weights supporting the same average propagation speed as the modulated chain, timing correction did not occur – spike pulses initiated 40 ms before or after a correct pulse remained about 40 ms behind or ahead (Fig 1C). Varying spike pulse initiation time across several hundred milliseconds confirmed that the temporal basin of error correction matched the 100 ms period of the external input (Fig 1D), while spike pulse timing in the unmodulated chain remained uncorrected regardless of start time (Fig 1E). One-dimensional but appropriately shaped external input can thus cause the network dynamics to follow the temporal structure of the input signal, while their sequential ordering is determined by the network’s internal connectivity (the ordered chain links, here).

### Sequence evolution through a spatiotemporal speed landscape

To understand theoretically how timing correction, intrinsic speed, and external input interact we analyzed a simplified, more general model of sequence propagation. For an evolving sequence whose instantaneous position is given by *x*(*t*) (the position of the propagating spike pulse in our network model), let its speed be given by:

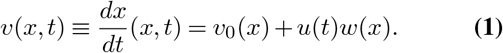

where *v*_0_(*x*), *u*(*t*), and *w*(*x*) are the intrinsic propagation speed profile, time-varying scalar input level, and spatial input weight profile. When *u*(*t*) = 0 (input is off) propagation speed depends only on *x* and equals the intrinsic speed *v*_0_(*x*). When *u*(*t*) = 1 (input on) the propagation speed evaluated at *x* is the sum of the intrinsic speed *v*_0_ and input weight *w* at *x*. Under what conditions does such a system cause sequences with perturbed start times to converge to the same timecourse?

To gain intuition we first examine how to stabilize the constant-speed sequence *x*(*t*) = *t* via the spatiotemporal “speed landscape” given in (1). Arbitrary sequences **z** = **f** (*t*) can be constructed by finding a map **z** = **g**(*x*) that transforms the *x* coordinate into an arbitrary value or vector (akin to activating sets of motor neurons (26)), so that the stability of **z**(*t*) follows the stability of *x*(*t*). As the input timecourse and spatial profiles of the intrinsic speed and input weights and are all 1-D, the speed landscape will have a rank-2 structure, the key bio-inspired constraint on the problem we seek to solve.

Fig 2 shows a temporal square-wave input *u*(*t*) oscillating between 0 and 1 and example *v*_0_(*x*) and *w*(*x*) that stabilize *x*(*t*) = *t*. The contributions of each term to the full speed landscape *v*(*x, t*) can be understood geometrically (Fig 2A-C). As the intrinsic speed *v*_0_(*x*) has no temporal dependence it admits variation only along the vertical *x* direction (Fig 2A), with certain horizontal bands corresponding to an intrinsic speed greater than 1 (purple) and others to an intrinsic speed less than 1 (orange) (Fig 2A). The input-modulation term *u*(*t*)*w*(*x*) admits both spatial and temporal variation, yielding a rank-1 contribution to the speed landscape (Fig 2B); certain regions correspond a positive modulation that increases propagation speed (purple) and others to a negative modulation that decreases propagation speed (orange). Summing the contributions of the intrinsic speed *v*_0_(*x*) (Fig 2A) and input modulation *u*(*t*)*w*(*x*) (Fig 2B) yields the full speed landscape *v*(*x, t*) under the fluctuating input (Fig 2C). In this example, *v*(*x, t*) = 1 can arise in regions with unity intrinsic speed and no modulation, regions with increased intrinsic speed but negative input modulation, or regions with decreased intrinsic speed but positive input modulation. Regions corresponding to *v*(*x, t*) *<* 1, which slow sequences down, or *v*(*x, t*) *>* 1, speeding sequences up, can each arise from either the intrinsic speed term *v*_0_(*x*) or the input modulation *u*(*t*)*w*(*x*). The temporal evolution of the output sequence is given by how *x*(*t*) moves through this landscape.

**Fig. 2.**
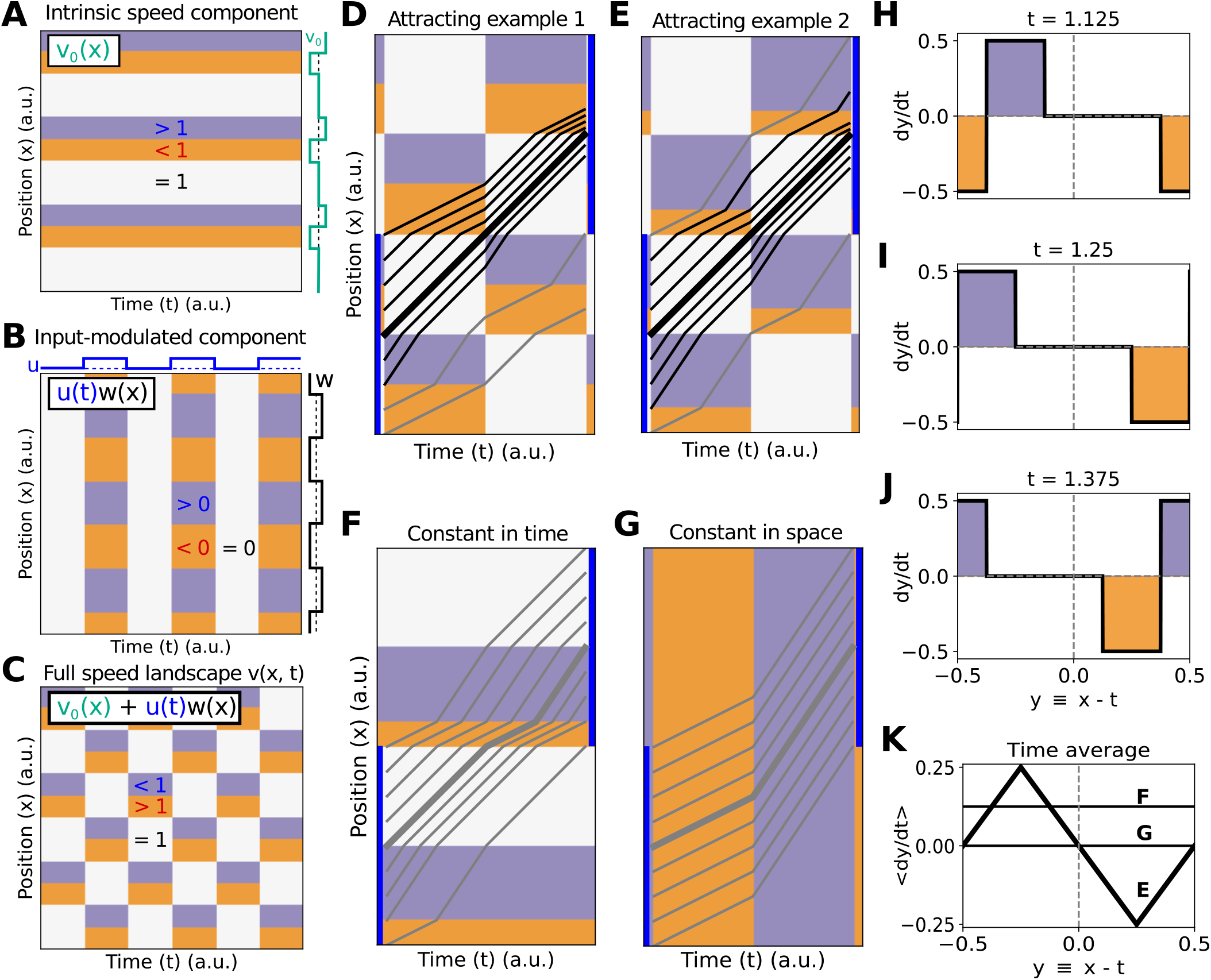
Time-locking attractors in example speed landscapes. A. Contribution of the position-dependent intrinsic speed profile to the speed landscape. Orange and purple indicate zones of slowed and hastened speeds, respectively. B. Contribution of spatiotemporal input modulation profile to the speed landscape. C. Full speed landscape with intrinsic speed and input modulation. Purple (fast) zones in input modulation profile cancel orange (slow) zones in intrinsic speed profile and vice versa, yielding a rank-2 checkerboard-like pattern. D. Multiple example sequences with different starting positions moving through speed landscape in C for one temporal period. Blue bars provide reference positional ranges offset by one spatial period. Black trajectories are within the basin of attraction. E. As in D but for a slightly different speed landscape. F. As in E but without temporal modulation. G. As in D/E/F but for a speed landscape without spatial modulation. H-J. Time-derivative of sequence position in a moving reference frame *y ≡ x − t* at different time points corresponding to E. K. Time-average of *dy/dt* over one temporal period for examples in E, F, and G.

Sequences that begin at different times or positions take different paths through the speed landscape. In our example in Fig 2 the sequence *x*(*t*) = *t* travels at constant speed, as desired, whereas sequences instantiated at slightly advanced or delayed positions pass through more slow or fast zones, respectively (Fig 2D). After a short time perturbed sequences will approach the sequence *x*(*t*) = *t* if the perturbation lies within a small enough window. This attractive effect in our example is robust to small changes in the speed landscape (Fig 2E), which only changes the basin of attraction. Thus the external input corrects, or stabilizes, the timing of perturbed sequences.

To identify formal conditions enabling timing correction we define a *time-locking attractor x*(*t*) = *t* as a stable fixed point in a constant-velocity moving reference frame. Letting the position coordinate in the moving frame be *y*(*t*) ≡ *x* − *t*, the function *dy/dt* = *f*_*y*_(*y*) describes the attraction or repulsion of the system toward the target sequence *x*(*t*) = *t*. Since *v*(*x, t*) is rank-2, *dy/dt* must also change over time because the local speed landscape surrounding *y* = 0 changes throughout the sequence (Fig 2H-J). In our example the time average ⟨*dy/dt*⟩ = ⟨*f*_*y*_(*y*) ⟩ has a downward zero-crossing at *y* = 0; thus, points *y <* 0 will hasten and points *y >* 0 will slow, on average, stabilizing *x*(*t*) = *t* and yielding the desired timing correction (Fig 2K). Crucially, for such a fixed point to exist, neither the input timecourse *u*(*t*) nor weight profile *w*(*x*) can be constant. When either is constant, all sequences spend the same total time in either slow or fast zones and thus have the same average speed and do not converge (Fig 2F, G); ⟨*dy/dt*⟩ lacks a downward-crossing zero at *y* = 0 so no correction occurs (Fig 2K). Thus, a scalar-valued external input can correct timing errors only if it fluctuates in both time and space. As shown in Fig 2-Supp-fig-1 and Fig 2-Supp-fig-2, time-locking attractors can also emerge under constant intrinsic speed and either purely excitatory or purely inhibitory modulation (reflecting the possible case that Uva provides a single sign of modulation onto HVC), so long as this modulation still fluctuates appropriately in both time and space. The resulting stable trajectories are more complex, however, since their propagation speed is not constant.

### Conditions for time-locking attractors

For a time-locking attractor to emerge generally, we found that it suffices for the relationship among the input timecourse and spatial profiles of the input weights and intrinsic speed to satisfy two conditions. First, the propagation speed *v*(*x, t*) should equal 1 when *x*(*t*) = *t*: using (1) we require

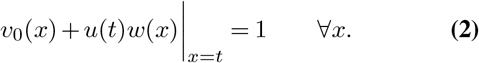

This condition creates a fixed point ⟨ *dy/dt*⟩ = 0 corresponding to *x*(*t*) = *t*. Differentiating (2) with respect to time yields

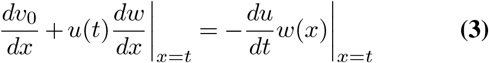

since *dx/dt* = 1.

For *x*(*t*) = *t* to be stable, we require from (1) that

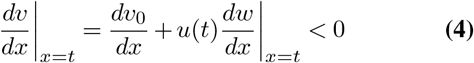

so that lagging trajectories *x*(*t*) = *t − δ* experience an increased propagation speed, whereas advanced trajectories *x*(*t*) = *t* + *δ* experience a decrease. Substituting (3) into (4) yields the sufficient condition

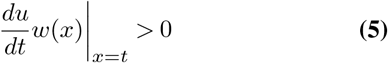

which requires *w*(*x*) to have the same sign as *u*(*t*) evaluated at *t* = *x*. Thus, when *u*(*t*) is increasing, *w*(*x*) |_*x*=*t*_ should be positive, and when *u*(*t*) is decreasing *w*(*x*)|_*x*=*t*_ should be negative. A simple (although not unique) solution is to let *w*(*x*) be proportional to the time-derivative of *u*(*t*) evaluated at *t* = *x*, and to then solve (2) to find *v*_0_(*x*) so that the stable trajectory has a uniform speed.

Time-locking attractors thus emerge when the input weight profile *w*(*x*) “reflects” spatially (has the same sign as) the input’s time derivative. As an input *u*(*t*) is not likely to continually increase for an arbitrarily long period in a real system, but will instead fluctuate in time, its spatial profile *w*(*x*) must accordingly fluctuate in space, although (5) implies that as long as the signs of *du/dt* and *w* match, their relationship need not be fine-tuned.

This rule allows us to construct input timecourses and weight profiles that propagate stable sequences. As an example, for *u*(*t*) = *cos*(*t*) + 1 we can use (5) to choose *w*(*x*) = −*sin*(*x*) and (2) to choose *v*_0_(*x*) = 1 +(*cos*(*x*) + 1)*sin*(*x*). Sequences initiated across several start times and positions reveal that these *u*(*t*), *w*(*x*), and *v*_0_(*x*) indeed yield a time-locking attractor *x*(*t*) = *t* (Fig 3A). Periodicity is not required, however. Choosing *w*(*x*) and *v*_0_(*x*) via the above rule when *u*(*t*) was a sample from a smoothed noise process also yielded the attractor *x*(*t*) = *t* (Fig 3B). Plotting *dy/dt* in the moving reference frame with *y* = *x* −*t* reveals that although the momentary phase portrait in each example is highly variable, its time average ⟨*dy/dt*⟩ yields a 1-D flow with a fixed point at *y* = 0 (Fig 3C-F), thereby stabilizing *x*(*t*) = *t*. Thus, given an arbitrary fluctuating 1-D input, there exist accompanying spatial profiles of the input weights and intrinsic speed that stabilize the evolution of the sequence *x*(*t*) = *t*, with the local temporal basin of attraction depending on the timescale of the input fluctuations.

**Fig. 3.**
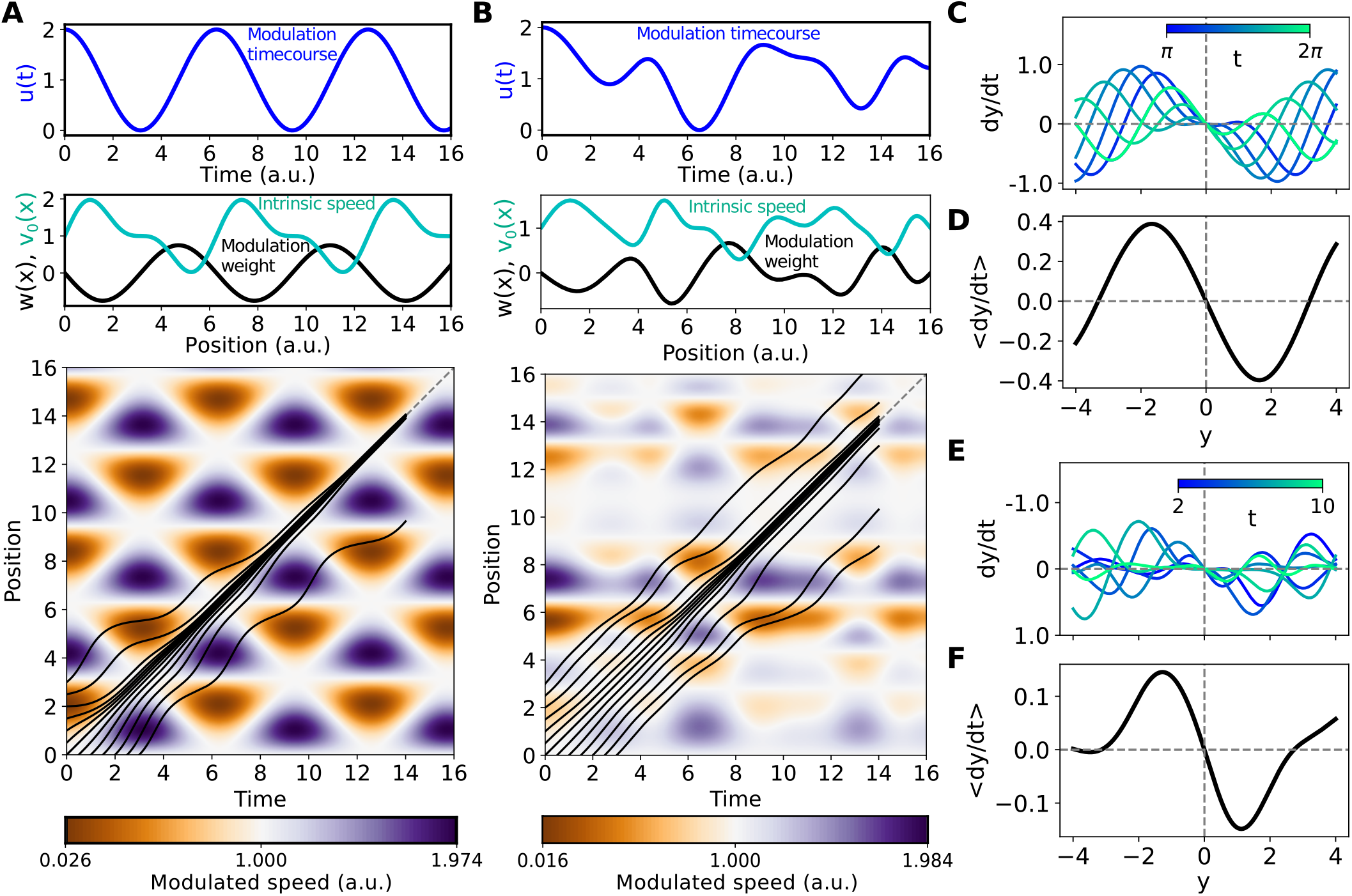
General conditions for a time-locking attractor to emerge from 1-D input. A. Example intrinsic speed and modulation weight profile (middle) derived from a periodic input timecourse (top), with *w*(*x*) *∝ du/dt*|*t*=_*x*_. Examples of sequence evolution through modulated speed landscape for various start times and positions (bottom). B. As in C but for an aperiodic input timecourse. C. Phase portrait of A in the moving reference frame *y = x −t* evaluated at several timepoints. D. Time average of momentary phase portraits in C (corresponding to speed landscape in A). E. Phase portrait of B in the reference frame *y* = *x − t* at several timepoints. F. Time averaged of phase portraits in E (corresponding to speed landscape in B). The time-averaged dynamics in D and F each exhibit a stable fixed point at *y* = 0 corresponding to the sequence *x*(*t*) = *t*.

### Correlated fluctuations in motor output timing

Due to the timing correction our model imposes, in the face of noise we expect durations of adjacent segments of the output sequence near the timescale of the external input to be anti-correlated. Segments shortened by noise will tend to be followed by segments slowed by the input’s correction effect, and vice versa for segments dilated by noise. We verified this by generating noisy sequences over the range *x* ∈ [0, 1] atop an input-driven speed landscape according to

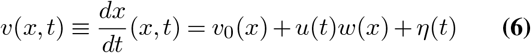

where *η*(*t*) is an Ornstein-Uhlenbeck process (exponentially filtered Gaussian white noise) with timescale *τ*_*η*_ = 1*/*30. Fig 4A shows example output sequences modulated by a periodic speed landscape. Whereas unmodulated sequences (*v*(*x, t*) = 1 + *η*(*t*)) diffuse freely, modulation by the input-dependent speed landscape constrains the spread (Fig 4A,B).

**Fig. 4.**
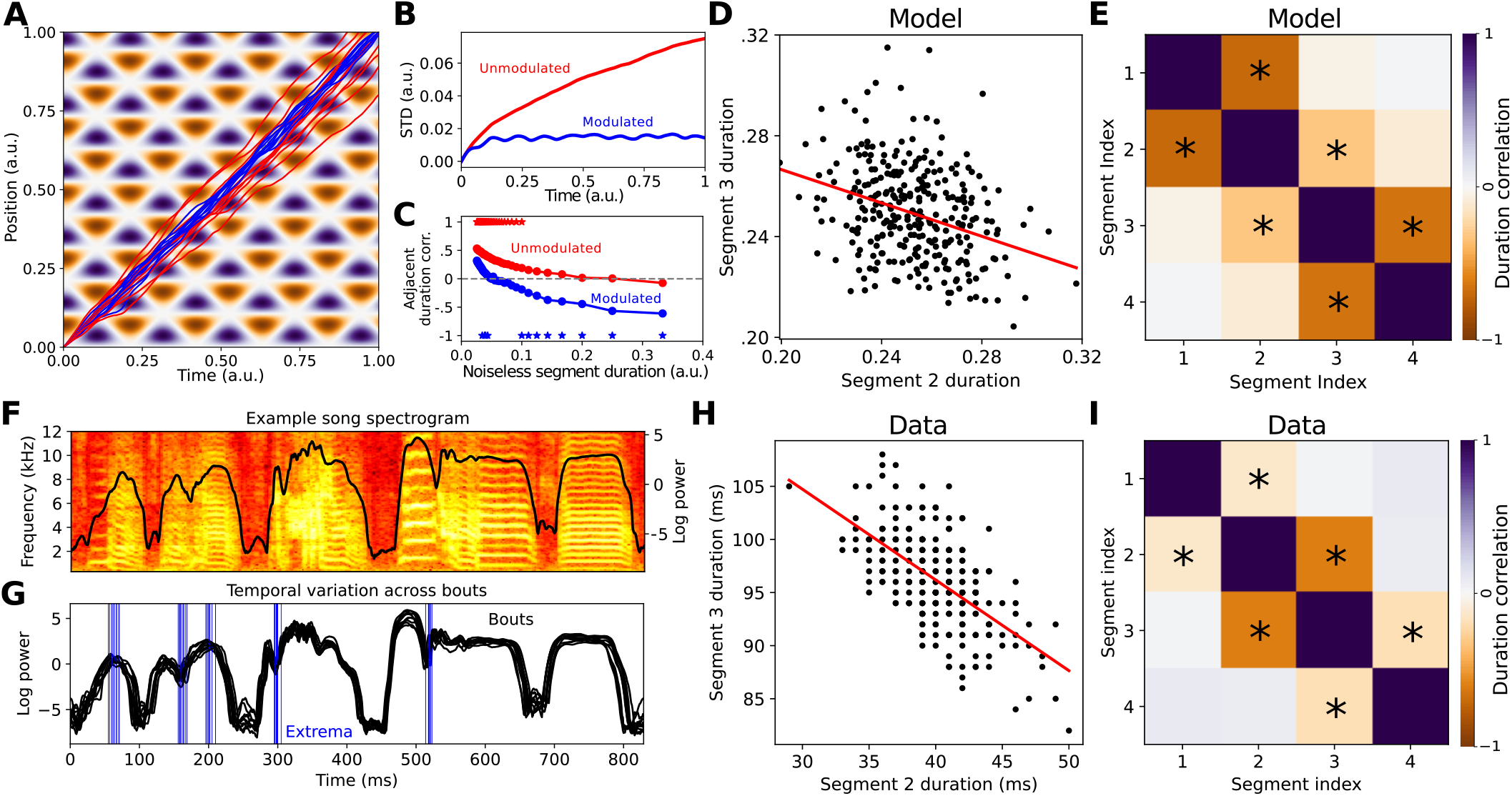
Correlated fluctuations in motor output timing. A. Example noisily propagating output sequences while modulated (blue) or not modulated (red) by underlying speed landscape (background) generated by 1-D input fluctuations. B. Growth of standard deviation of sequence position over time for modulated and unmodulated sequences, corresponding to 300 sample sequences as demonstrated in A. C. Pearson correlation (across trials) between durations of adjacent segments (averaged across segment pairs) of the output sequence vs the noiseless segment duration, for modulated (black) and unmodulated (orange) trials. Stars indicate an average *p <* .05 (two-sided t-test for zero-valued correlation coefficient) across segment pairs. D. Durations of second and third segments (given 4 equal segments total, each with a noiseless duration of .25) across trials, and best fit line (*R* = .324, *p <* 10^*−*8^). E. Correlation matrix between durations of all segment pairs in simulated motor output. Stars indicate correlations with *p <* .05. F. Spectrogram of example zebra finch song motif, with log power overlaid in black. G. Log power of the same song motif across several bout renditions in one bird. Blue lines show extrema used to define song segment boundaries before quantifying variation across renditions. H, I. As in D, E but for real zebra finch song segments (N = 292). (R = -.62, p < 10^*−*31^ for best fit line in H.)

To investigate temporal structure in the output sequences we split the output into even segments and examined the durations over which they unfolded. Durations of sufficiently short adjacent segments (noiseless duration ≪ *τ*_*η*_) were positively correlated, regardless of modulation by the speed landscape, as they were likely to receive similar noise. In the input-modulated sequences, however, sufficiently long segments became anti-correlated, a sign of timing correction that was not present without the modulation (Fig 4C-E).

We tested our model’s prediction that durations of certain segments of motor output sequences should be anti-correlated by analyzing real zebra finch songs. Songs were produced by adult males in isolation (“undirected” song), with the recorded audio converted to spectrograms using a short-time Fourier transform (Fig 4F). Due to the imprecision in determining exactly when an arbitrary moment *x* in a song occurred (and spurious anti-correlations introduced by jitter in such a labeling process), instead of splitting the song into even segments we examined segments bounded by peaks and valleys in the power of the spectrogram, retaining only segment boundaries that could be reliably identified in every song (Fig 4F-G). Indeed, durations of adjacent segments of zebra finch song were anti-correlated (Fig 4H-I), as predicted by and dependent on the input-modulation in our model, a finding that held up across multiple birds (Fig 4-Supp-fig-1, A-H). This suggests the neural song circuit in songbirds may recruit a similar mechanism for timing control as the input-dependent time-locking attractors we have investigated.

## Discussion

We introduced a model of precision motor timing via scalar input fluctuations and validated a key prediction in recorded zebra finch songs. When a scalar input locally and dynamically controls how fast a sequence unfolds and the input’s spatial structure reflects its time derivative, time-locking attractors emerge that continually correct timing errors without requiring knowledge of the errors. Although timing errors could accumulate in the input itself, they do not additionally accumulate in the downstream sequence generator. This allows multiple motor processes receiving the same input fluctuations to stay coordinated for long periods, reflecting how left and right HVC may receive similar input from left and right Uva. Our work builds on previous observations of spatiotemporal inhomogeneities in songbird HVC (27), providing new insight into how spatial and temporal signals may be linked to temporally coordinate ongoing neural activity.

While neural spike time coordination has been explored at length in “synfire” chain models (1, 3, 24, 28, 29), where synchrony arises through extensive excitatory coupling among neurons between network layers, our work addresses the distinct problem of quenching timing errors via external input. For instance, HVC-inspired computational work showed that zones of inhibitory feedback triggered by a propagating spike pulse could keep the spikes within the pulse synchronized without redundant excitatory connections (29). Our model expands on this picture by explaining how a propagating spike pulse or other sequential activity pattern could be kept synchronized with an external input, and in turn with other sequential activity patterns receiving the same input, all without requiring feedback about timing errors. A separate input-based model of neural timing showed how external oscillatory inputs matched to spike latencies in a feed-forward network could entrain the propagation of a spike pulse by imposing specific windows at which spikes can occur (16), suggestive of hippocampal spike sequences that are locked to high-frequency “ripple” events (30). Our work presents a distinct input-based mechanism grounded in speed control rather than neuronal properties (although still applicable to spiking networks [Fig 1]) and which requires time-varying but not necessarily oscillating input.

Our model effectively splits each motor process into a dynamical system specifying sequence order and a scalar input controlling timing. This is a simple and perhaps biologically favorable alternative to directly coupling multiple motor networks, which for long sequences would require coupling at many points over the sequence. During the course of development allocating timing precision to an external input could allow other features of the output sequence, such as syntax or instantaneous spectral content (19, 31) to be learned independently of timing, which might simplify and hasten learning.

How might the match between our input’s time-derivative *du/dt* and spatial profile *w*(*x*) emerge or be maintained over time, for instance in the face of random synaptic fluctuations (32–34)? Recalling that *w*(*x*) in our spiking network model (Fig 1) represents the synaptic weight from the external input onto the neurons at position *x* in the chain, the relationship between *du/dt* and *w*(*x*) could potentially be stabilized by local spike-timing-dependent synaptic plasticity (STDP) rules (e.g. (35–39)). For instance, to keep the input synaptic weight *w*(*x*) sufficiently positive when *du/dt* _*t*=*x*_ is positive (following the constraint in Eqn. 5), it could suffice to increase *w*(*x*) whenever a spike pulse passing through the chain link at position *x* at time *t* is accompanied by accelerating presynaptic spikes from the external input onto that chain link at *t*. One average, one might expect such a scenario to yield more postsynaptic-followed-by-presynaptic spike pairs–if the presynaptic spike rate is increasing, then in a small window around *t* there will be more presynaptic spikes in the later part of the window. An anti-Hebbian STDP rule (38) might then suffice to increase *w*(*x*) in response to the repeated occurrence of these spike pairs. Conversely, when *du/dt*|_*t*=*x*_ is negative, so that *w*(*x*) should also be sufficiently negative to satisfy (5), one might expect more presynaptic-followed-by-postsynaptic spike pairs. If *w*(*x*) were represented by an inhibitory synapse, a Hebbian STDP rule (35, 36, 39) could increase the weight of that synapse to make *w*(*x*) more negative. While more work will be required to illuminate the details of how such plasticity mechanisms could stabilize or even give rise to the spatial input profile suggested by our model, the local nature of the plasticity required along with the lack of fine-tuning in our model suggest a bioplausible maintenance or development process that could achieve this.

We expect the role of thalamocortical inputs we have proposed to co-exist with other thalamic functions and multiarea interactions. Baseline Uva input to HVC, for instance (21), may be required for song production regardless of temporal precision; thalamic inputs also likely contribute to setting movement preparatory states in cortex (12–15). Thus, one may expect to observe multiple functions superimposed in thalamocortical input dynamics measured experimentally. In birds Uva and HVC are embedded within a larger brainstem-thalamocortical loop (22) that indirectly couples the two hemispheres, which may be involved in the slowing of song when HVC is only cooled unilaterally (40). Our model posits that such coupling is nonetheless strictly gated by low-dimensional Uva inputs to HVC, with Uva sending predominantly timing signals, while all ordering and spectral information is contained within HVC’s recurrent and downstream connectivity (although other HVC inputs may contain additional syntactic information, i.e. influencing the order in which HVC neurons fire (19)). Our work thus contrasts with models of birdsong neurophysiology in which the network connections encoding sequence order are coiled around the brainstem-thalamocortical loop (41, 42); we instead ascribe sequence ordering and timing functions predominantly to HVC and Uva, respectively.

Although not easily dissociated from the multi-functional system it is embedded in, our model makes additional predictions. First, it predicts that Uva inputs onto HVC should exhibit significant spatial structure relative to song position. Specifically, HVC neurons that spike during Uva increases should receive more excitation from Uva, and those that spike during decreases more inhibition (possibly via HVC interneurons). Although technically challenging, one could test this by tracing Uva-HVC projections and recording which HVC neurons spike at different points during song. More generally, our model predicts that removing temporal fluctuations from thalamic input to motor cortex should preclude timing error correction. This could be tested by silencing thalamus and directly activating thalamocortical axon terminals (10) with constant or ramping stimulation, which should increase variability in the total durations of the accompanying cortical and motor sequences, relative to stimulation matched to real thalamic fluctuations.

The central mechanism in our model applies to sequence generators beyond chain networks. One only requires a sufficiently ordered sequence to support a notion of timevarying position *x*(*t*), and a fluctuating input that modulates *dx/dt* differently at different *x*. The first condition is met, for instance, by neural activity that evolves along a timeordered manifold, a topology hypothesized to underlie dynamics in several cortical areas (43), or through a sequence of metastable network attractors (9). To model timing control in such a system one could extend work on input-dependent speed control of recurrent network dynamics (17) by making the input itself (presumably coming from another brain area) dynamic and localizing its slowing and/or accelerating effects to different regions of the sequence. Finally, time-locking attractors from scalar input fluctuations may have applications outside neuroscience, for instance in coordinating cellular processes via fluctuating molecular concentrations or synchronizing growth processes across a population of organisms via fluctuating temperature or sunlight.

## Methods

### Neuron and synaptic dynamics

Our spiking network used integrate-and-fire neurons, which receive conductance inputs from excitatory and inhibitory neurons, and in which adaptation is modeled as a self-inhibitory synaptic current:

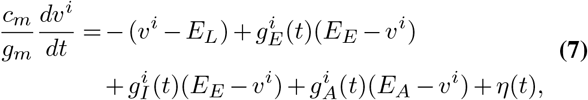

where

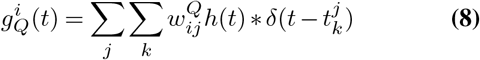

and *v*^*i*^ is the *i*-th neuron’s membrane voltage; *η* is a white noise current; *c*_*m*_ = 1*µF/cm*^2^ and *g*_*m*_ = 100*µS/cm*^2^ are the membrane leak capacitance and conductance, yielding a membrane time constant of *τ*_*m*_ = *c*_*m*_*/g*_*m*_ = 10ms; *E*_*L*_ = −60*mV, E*_*E*_ = 0*mV, E*_*I*_ = −80*mV, E*_*A*_ = −100*mV* are the leak, excitatory, inhibitory, and adaptation reversal potentials; 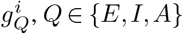 are the relative excitatory, inhibitory, and adaptation conductances; 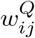 is the weight of synapse type *Q* onto neuron *i* from *j*; 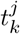 is the *k*-th spike time of neuron *j*; and *h*(*t*) is an exponential filter with time constants *τ*_*E*_ = *τ*_*I*_ = 2*ms, τ*_*A*_ = 10*ms*. Neurons spiked when *v*^*i*^ ≥ *v*_*th*_ = −50*mV* and were reset to *E*_*L*_ = −60*mV* for a 2*ms* refractory period. The simulation timestep was Δ*t* = 5*ms*, and *η*Δ*t ∼* N (0, *σ*^2^ = .01*nA*^2^*s*).

### Network architecture

Our spiking network comprised 321 chain “links”, each containing 30 excitatory neurons. Recurrent (within a link) weights were only excitatory and equal to 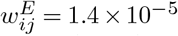 with probability .6 and 0 otherwise. Feed-forward weights were excitatory and all-to-all from each link to its successor with 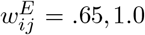, or 1.4 10^−5^, corresponding to weak, medium or strong connections (Fig 1B). Adaptation “weights” for all neurons were 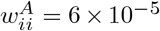. All feed-forward weights being set to “medium” supported a self-sustained spike burst (approx. 3-5 spikes over 5-10 ms in each neuron) propagating from link to link. Last, neurons received E and I synapses capable of transmitting Uva-like input spikes, with 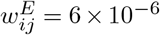 and 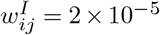 for E or I external inputs. In Fig 1B, contiguous sets of 58 chain links alternated between receiving external E or I input synapses, and feed-forward weights alternated between 58 medium weights, 29 weak weights, 29 strong weights, etc. (black). In Fig 1C, 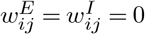 for all external inputs, and all feed-forward weights were medium. In Fig 1B, external inputs alternated between a 50 ms “on” state and a 50 ms “off” state corresponding to 800 Hz and 0 Hz input spike frequencies, and in Fig 1C were off throughout the whole simulation.

### Time-locking attractor simulations

Example output sequences in Fig 2D-G began at 9 evenly spaced start positions spanning one spatial period of the shown speed landscape. In Fig 3, simulations used an integration timestep of Δ*t* = .001. In Fig 3D, *u*(*t*) was sampled from a white-noise process smoothed with a Gaussian kernel with width .9*/*Δ*t*; *w*(*x*) was chosen to be .65*du/dt*|_*t*=*x*_, and *v*_0_(*x*) = 1 − *u*(*x*)*w*(*x*).

### Data analysis

Song renditions were recorded from adult male zebra finches during an “undirected” song context (no female present) and transformed into spectrograms using a short-time Fourier transform. The power of the logarithm of the spectrogram was computed at each timepoint to get a scalar representation of song, from which peaks were identified and used to define song segment boundaries (only peaks that could be reliably identified across renditions were used as boundaries). Durations of each segment were then extracted and correlations among resulting segment durations computed across song renditions.

## Acknowledgments

We would like to acknowledge Nader Nikbakht and Jonathan Pillow for helpful discussions regarding the preparation of this manuscript. This work was supported by the Simons Foundation’s Simons Collaboration for the Global Brain and by NIH 1R01NS104925.

## Notes

### Competing Interest Statement

The authors have declared no competing interest.

