## Supplementary Figures for "Precision motor timing via scalar input fluctuations"

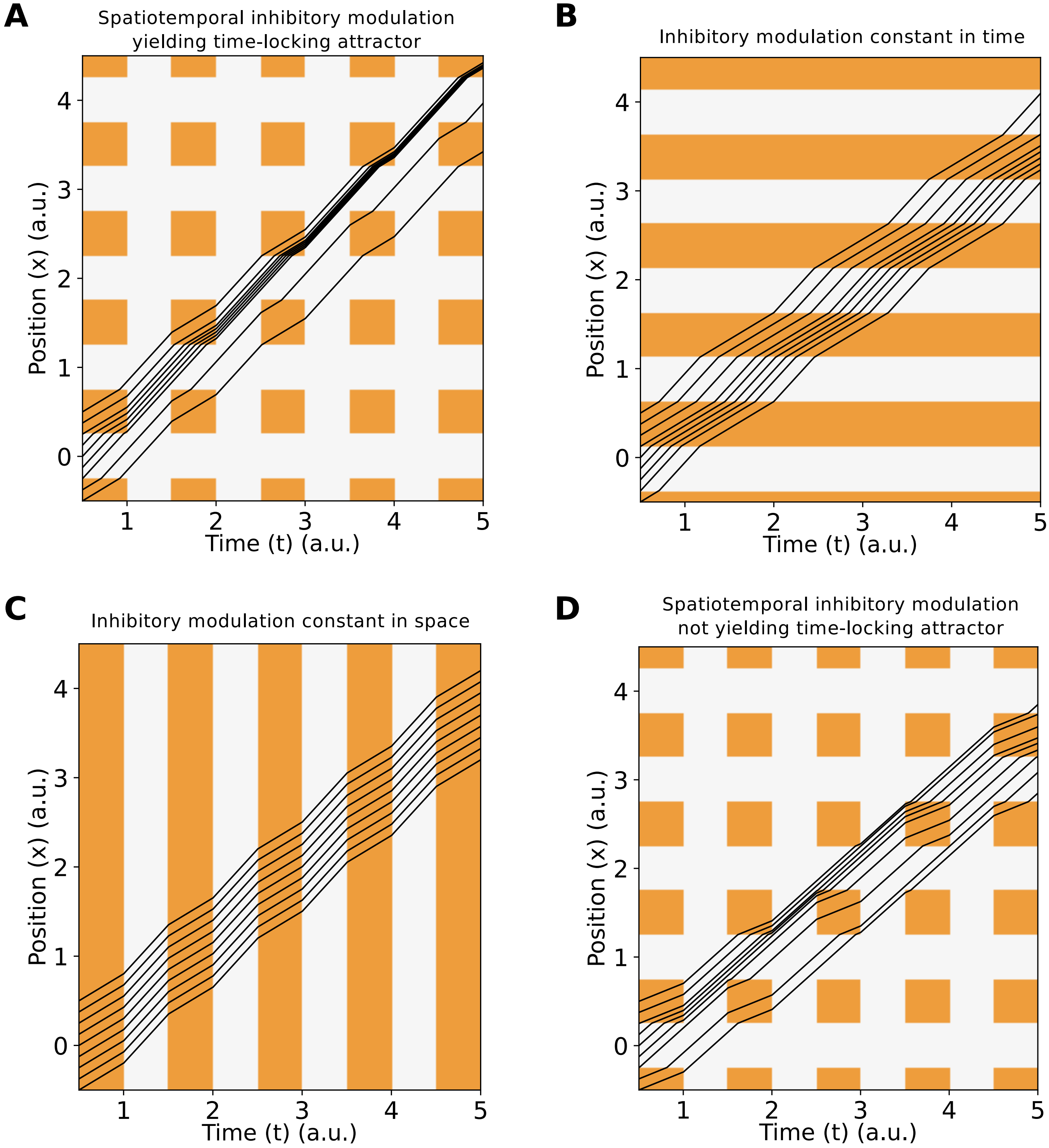


**Fig 2-Supp-1: Example speed landscapes using purely inhibitory modulation**. Black traces show trajectories beginning at different starting points. Orange zones indicate spatiotemporal regions in which propagation speed slows relative to the unmodulated intrinsic speed (white). A. Speed landscape where modulation timecourse and spatial profile are matched to yield time-locking attractor. B. Speed landscape from modulation constant in time. C. Speed landscape from modulation constant in space. D. Speed landscape with both temporally and spatially varying modulation but which does not yield a time-locking attractor. Here the intrinsic speed (slope of the black lines as they pass through white regions) is sufficiently lower than that of the landscape in A that no time-locking attractor emerges.

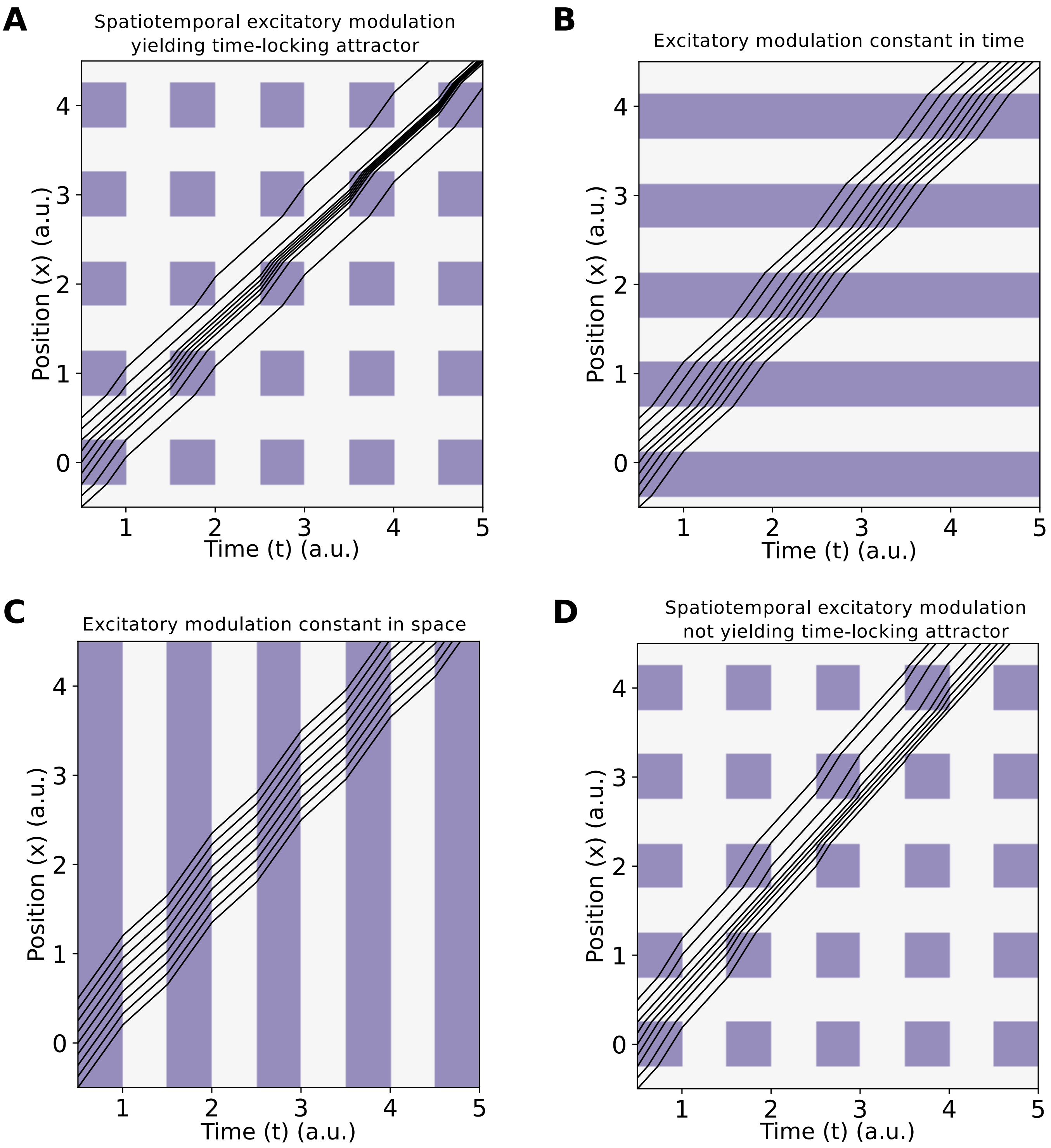


**Fig 2-Supp-2: Example speed landscapes using purely excitatory modulation.** As in Figure 2-Supp-1, except where speeds can only increase via modulation (purple zones). In D, the intrinsic speed is sufficiently higher than in A that no time-locking attractor emerges.

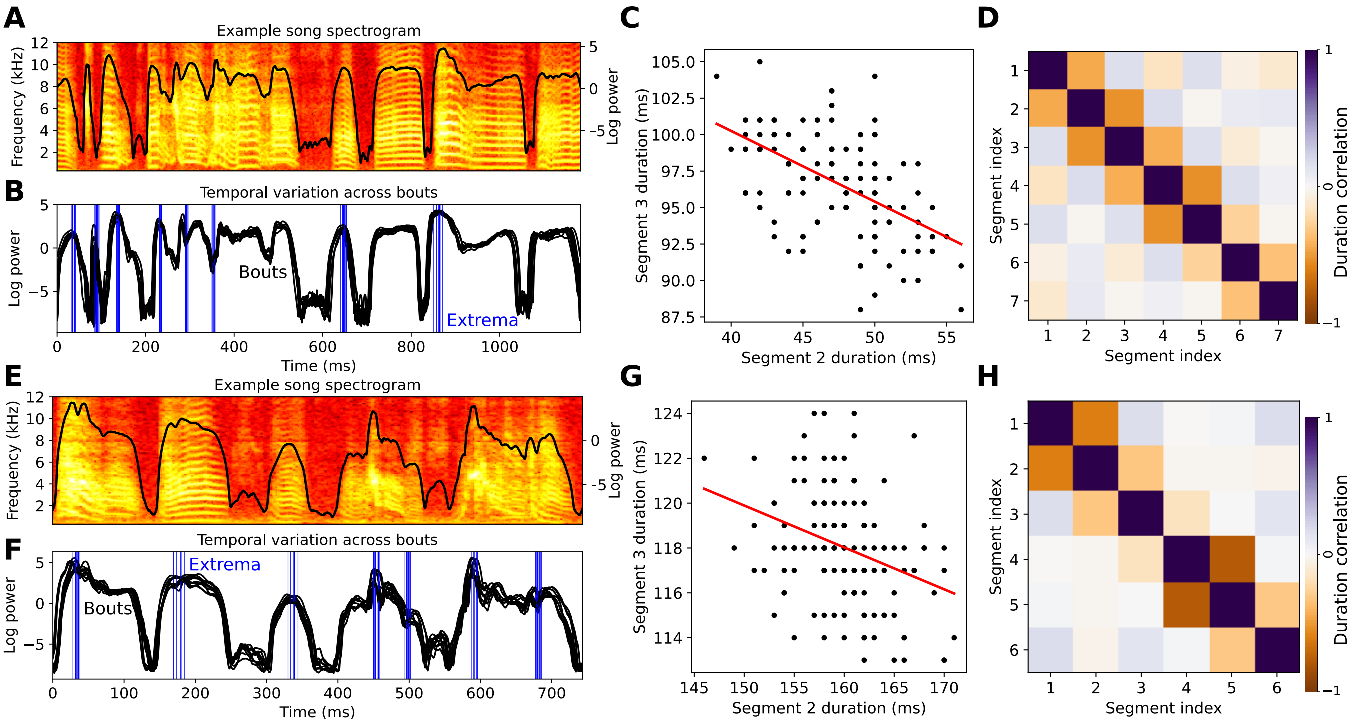


**Fig 4-Supp-1: Anti-correlated adjacent song segment durations in additional birds.** As in Fig 4F-I, but for two other birds. $R = -.566, p < 10^{-10}$ for best fit line in C; $R = -.326, p < 10^{-4}$ in G.
